# SCAMP3 is essential for proper formation and function of neutrophil granules

**DOI:** 10.1101/2025.08.15.670532

**Authors:** Almke Bader, Jincheng Gao, Nina Reiter, Devon Siemes, Annette Zehrer, Bettina Schmid, Ignasi Forné, Daniel Robert Engel, Barbara Walzog, Daniela Maier-Begandt

## Abstract

Host defense functions of neutrophils during infection critically depend on microbicidal and proteolytic proteins stored in primary, secondary and tertiary granules that are released into the phagosome or into the extracellular space upon degranulation. Granules are generated during granulopoiesis and impaired granule production or granule protein sorting has been linked to inefficient pathogen clearance resulting in recurrent bacterial and fungal infections. Here, we studied the role of the membrane protein secretory carrier associated membrane protein 3 (SCAMP3) for neutrophil defense functions. We generated *Scamp3* knockout (KO) Hoxb8 cells and found that killing of *Escherichia coli* by *Scamp3* KO Hoxb8 cell-derived neutrophils (dHoxb8 cells) was compromised as compared to control dHoxb8 cells *in vitro*. Mass spectrometric and Western blot analyses revealed a significant reduction of primary, secondary, and tertiary granule proteins in the genetic absence of *Scamp3*, resulting in a reduced overall granularity of these cells. Accordingly, degranulation was reduced in *Scamp3* KO dHoxb8 cells compared to control dHoxb8 cells. Similarly, SCAMP3 deficiency in zebrafish resulted in reduced neutrophil granularity in comparison to wild-type animals. However, neutrophil migration towards sites of *E. coli* infection was unaffected in *scamp3* KO zebrafish larvae. In summary, SCAMP3 represents an important novel player in granule equipment and degranulation, with key functions in neutrophil defense mechanisms during host-pathogen interactions *in vitro*.

**Brief summary sentence:** The membrane protein secretory carrier associated membrane protein 3 (SCAMP3) is essential for proper protein equipment of neutrophil granules and *Scamp3*-deficient neutrophils have an impaired bacterial killing capacity *in vitro*.

## 1 Introduction

Neutrophils are the most abundant leukocytes in the blood and the first cells to be recruited to sites of inflammation where they can eliminate cell debris or combat invading pathogens.^1–3^ Neutrophils play an essential role for host defense against invading pathogens and when their killing capacity is impaired, patients suffer from severe, life-threatening infections.^3,4^ Pathogen elimination is achieved by different mechanisms including phagocytosis, degranulation, production of reactive oxygen species and formation of neutrophil extracellular traps.^2^

During degranulation, microbicidal and proteolytic proteins are released into the extracellular space or into the phagosome.^5^ Neutrophils are equipped with a heterogeneous population of granules that can be distinguished into three different types, namely primary (azurophil), secondary (specific) and tertiary (gelatinase) granules. Additionally, neutrophils contain secretory vesicles. Generation of granules and secretory vesicles takes place during granulopoiesis where primary granules are formed at the stage of promyelocytes, secondary granules at the stage of myelocytes and tertiary granules at the stage of metamyelocytes and band cells. Lastly, secretory vesicles are formed via endocytosis at the stage of band cells and mature neutrophils.^5^

According to the targeting-by-timing hypothesis, equipment of granules with their subset of proteins occurs by constantly rerouting newly build proteins from the endoplasmic reticulum-Golgi pathway into the granules without specific sorting into granule subsets.^6^ Thus, each type of granule contains overlapping but also specific marker proteins. Myeloperoxidase (MPO), neutrophil elastase (NE) and cathepsin D, amongst others, are specifically stored in primary granules.^5,7,8^ Lactoferrin (LTF), neutrophil gelatinase-associated lipocalin (NGAL) and olfactomedin 4 (OLFM4) are specifically located in secondary granules, and neutrophil collagenase (MMP8), matrix metalloproteinase-9 (MMP9) and lysozyme C-2 (LYZ2) are mainly found in tertiary granules.^5,7,9^ Despite extensive research on granule formation, the precise molecular mechanisms underlying the generation of granules and sorting of various proteins into granules remain incompletely understood.

The membrane protein secretory carrier associated membrane protein 3 (SCAMP3) belongs to the SCAMP family consisting of SCAMP1-5 with SCAMP1-4 being ubiquitously expressed and SCAMP5 mainly in neuronal cells.^10,11^ SCAMP3 harbors four transmembrane domains, a proline rich region that enables protein-protein interactions, with an intracellularly located N- and C-terminus. SCAMP3 is localized in almost all membranes of the cell but especially in the *trans* Golgi network and the endosomal system, and regulates the localization, recycling and degradation of proteins within the cell.^12–15^ In recent years, SCAMP3 has also been shown to play a role during bacterial and viral infections. Here, *Salmonella enterica* redistributes SCAMP3 from the *trans*-Golgi network to form *Salmonella*-induced SCAMP3 tubules and uses SCAMP3 for clustering and central localization of *Salmonella*-containing vacuoles.^16^ Similarly, different viruses such as rabies virus or enteroviruses interact with SCAMP3 to enhance viral replication.^17,18^ Interestingly, SCAMP3 was found to be localized in secretory vesicles, secondary and tertiary but not primary granules of human neutrophils.^19^ However, the functional role of SCAMP3 in neutrophils has not been elucidated yet. Hence, we set out to decipher the role of SCAMP3 for neutrophil functions. Using the murine Hoxb8 cell system, we found that *Scamp3* deficiency resulted in impaired bacterial clearance in *Scamp3* knockout (KO) Hoxb8 cell-derived neutrophils (dHoxb8 cells) compared to *Scamp3* control (CTRL) dHoxb8 cells infected with *Escherichia coli*. The absence of SCAMP3 led to reduced neutrophil granularity and reduced amounts of granule proteins. We confirmed the reduced granularity in neutrophils isolated from *scamp3*-deficient adult zebrafish. However, *scamp3* was dispensable for neutrophil migration to sites of infection and bacterial clearance *in vivo*. Thus, SCAMP3 may represent a novel regulator of granule formation and function, enabling efficient bacterial killing of neutrophils *in vitro*.

## 2 Material and Methods

### 2.1 Cell culture

Hoxb8-SCF cells (Hoxb8 cells) were generated and cultured as described previously^20,21^ in RPMI 1640 (Sigma-Aldrich) supplemented with 10% FCS (Sigma-Aldrich), 100 U/ml penicillin/100 μg/ml streptomycin (Sigma-Aldrich), 4% SCF-containing CHO cell supernatant, 30 µM β-mercaptoethanol (AppliChem) and 1 µM β-estradiol (Sigma-Aldrich). SCF-producing CHO cells and the plasmids pMSCVneo-ER-Hoxb8 and pCL-Eco for generation of Hoxb8 cells were kindly provided by Hans Häcker (School of Medicine, University of Utah). For differentiation towards neutrophils, β-estradiol was removed and 20 ng/ml recombinant murine G-CSF (PeproTech) was added to the medium for 4 days. Hoxb8 cells were kept at 37 °C, 5% CO_2_, and 95% humidity.

### 2.2 Generation of Scamp3 knock-out Hoxb8 cells

*Scamp3* knock-out Hoxb8 cells were generated by clustered regularly interspaced short palindromic repeats (CRISPR)/Cas9 technique. The guide RNA (gRNA, GACTCCGGCGAGCTTGACAA) was cloned into the lentiCRISPR v2 vector. HEK293T/17 cells (CRL-11268, American Type Culture Collection) were transfected with this lentiCRISPR v2-gRNA vector, psPAX2, and pCMV-VSV-G using Lipofectamine 2000 (Invitrogen) to generate virus particles. The virus-containing supernatant was filtered through a 0.45 µm filter, and used to transduce wild-type Hoxb8 cells with spinoculation (1000 x g, 90 min) in presence of Lipofectamine (Invitrogen). Transduced cells were selected with puromycin, subcloned, and screened for mutations in the *Scamp3* gene using the following primers for genotyping: 5’-GCTTAACATGAAGCTACCTGGG-3’ (forward) and 5’-CACCTTCACATAGCGACACTTC-3’ (reverse). Two cell lines with a homozygous knock-out of the *Scamp3* gene were selected (KO 01 and 02) and one cell line which underwent the CRISPR/Cas9 procedure but without induction of a mutation in the *Scamp3* gene was selected as control (CTRL).

LentiCRISPR v2 vector was a gift from Feng Zhang (Addgene plasmid # 52961; http://n2t.net/addgene:52961; RRID:Addgene_52961^22^). psPAX2 was a gift from Didier Trono (Addgene plasmid # 12260; http://n2t.net/addgene:12260; RRID:Addgene_1226). pCMV-VSV-G was a gift from Bob Weinberg (Addgene plasmid # 8454; http://n2t.net/addgene:8454; RRID:Addgene_8454^23^).

### 2.3 Flow cytometry

For flow cytometric analyses of cell surface protein expression, 5 x 10^5^ Hoxb8 cells at day 0 or day 4 of differentiation were collected and incubated with the following antibodies in PBS containing 5% FCS for 25 min at 4 °C: PerCP-eFluor710 anti-CD117 (1:200, clone 2B8, 46-1171-82, Invitrogen), eFluor660 anti-CD34 (1:50, clone RAM34, 50-0341-82, Invitrogen), PerCP-eFluor710 anti-Ly6G (1:100, clone 1A8-Ly6G, 46-9668-82, Invitrogen), AlexaFluor647 anti-CXCR2 (1:100, clone SA044G4, 149306, BioLegend), and the appropriate isotype controls. Cells were subsequently washed, resuspended in PBS containing 5% FCS, and analyzed using a CytoFLEX S flow cytometer (Beckman Coulter). Analysis was performed with FlowJo^TM^ software (BD Biosciences).

Granularity of unlabeled, differentiated Hoxb8 cells was assessed through the sideward scatter using a CytoFLEX S flow cytometer (Beckman Coulter) with subsequent data analysis with FlowJo^TM^ software (BD Biosciences).

### 2.4 Mass spectrometry

For whole proteome mass spectrometric analyses of *Scamp3* CTRL and KO cell lines, 1 x10^6^ cells on day 0 and day 4 of differentiation were collected by centrifugation at 300 x g for 5 min and the cell pellets were immediately frozen at -80 °C. Proteins from the different cell lines were digested using the iST kit (Preomics) following the recommendations of the manufacturer. For LC-MS/MS purposes, desalted peptides were injected in an Ultimate 3000 RSLCnano system (Thermo Fisher Scientific), separated in a 25 cm analytical column (Odyssey column, 75µm x 25cm, 1.7µm, 120 Å, IonOpticks) with a 90 min gradient from 4 to 40% acetonitrile in 0.1% formic acid. The effluent from the HPLC was directly electrosprayed into a Qexactive HF (Thermo Fisher Scientific) operated in data dependent mode to automatically switch between full scan MS and MS/MS acquisition. Survey full scan MS spectra (from m/z 375–1600) were acquired with resolution R=60,000 at m/z 400 (AGC target of 3 x 10^6^). The 10 most intense peptide ions with charge states between 2 and 5 were sequentially isolated to a target value of 1 x 10^5^, and fragmented at 27% normalized collision energy. Typical mass spectrometric conditions were: spray voltage, 1.5 kV; no sheath and auxiliary gas flow; heated capillary temperature, 250°C; ion selection threshold, 33.000 counts.

MaxQuant 2.1.0.0 was used to identify proteins and quantify by LFQ with the following parameters: Database, UP000000589_10090_Mmusculus_20210405.fasta; MS tol, 10ppm; MS/MS tol, 20ppm Da; Peptide FDR, 0.1; Protein FDR, 0.01 Min. peptide Length, 7; Variable modifications, Oxidation (M); Fixed modifications, Carbamidomethyl (C); Peptides for protein quantitation, razor and unique; Min. peptides, 1; Min. ratio count, 2. Missing abundances were imputed using a column wise random uniform 5th-10th percentile strategy. Gene set enrichment analysis was generated using GSEA (v.4.3.3) from MSigDB^24^ with the gene set databases M5:GO BP & CC (v2024.1.Mm) and enrichment statistic set to classic. Proteins were sorted by signal-to-noise ratio (SNR) which was determined using SNR = (<u>^-x**-**^</u>^1^<u>^---x**-**z-^</u>) with x-and x---being the mean values and d1 and d2 as the respective standard deviation. The enrichment dotplot was generated with ggplot2^25^ in R.

### 2.5 Bacterial killing assay

For analysis of bacterial uptake and survival, *Escherichia coli* MG1655 carrying the plasmid pEB2-E2-Crimson were used. pEB2-E2-Crimson was a gift from Philippe Cluzel (Addgene plasmid # 104010; http://n2t.net/addgene:104010; RRID:Addgene_104010^26^). *E. coli* were grown in LB medium (Carl Roth) containing 50 µg/ml kanamycin (Sigma-Aldrich) over night at 37 °C shaking at 180 rpm and were diluted to an OD_600_ of 0.01 and cultured for another 2 to 2.5 h at 37 °C shaking at 180 rpm. Per sample 2 x 10^5^ differentiated Hoxb8 cells were collected and resuspended in PBS containing 10% FCS. Where indicated cells were pre-treated with 10 µg/ml cytochalasin D for 15 min at 37 °C to prevent phagocytosis. *E. coli* were added to the cells with a multiplicity of infection (MOI) of 10, with an OD_600_ of 1 corresponding to 5.4 x 10^8^ bacteria/ml, and incubated for 30 min at 37 °C. Then cells were centrifuged, resuspended in 100 µg/ml gentamicin (Sigma-Aldrich) in PBS containing 10% FCS, and incubated for 1 h at 37 °C to kill extracellular bacteria. Subsequently, cells were washed and lysed with 0.1% saponin in PBS for 5 min at room temperature. Lysates were plated in appropriate dilutions on LB agar (Carl Roth) plates containing 50 µg/ml kanamycin and incubated at 37 °C over night. *E. coli* colonies were counted manually.

### 2.6 Degranulation assay

For analysis of degranulation, 2 x 10^5^ differentiated Hoxb8 cells were collected by centrifugation and resuspended in adhesion medium (Hank’s balanced salt solution (Bio West) containing 20 mM HEPES (AppliChem), 0.25% BSA (Sigma-Aldrich), 0.1% glucose (Merck Milipore), 1.2 mM Ca^2+^ (AppliChem), and 1 mM Mg^2+^(AppliChem)). Cells were incubated with 0.15% DMSO (AppliChem) as control or 10 µM fMLP (Sigma-Aldrich) and 10 µg/ml cytochalasin D (AppliChem) for 10 min at 37 °C. After centrifugation at 4 °C, the supernatant was collected and immediately frozen at -80 °C. Commercially available ELISA kits detecting MPO (HK210, Hycult Biotech), lactoferrin (G-MOES01235.96, Assay Genie), or MMP9 (MMPT90, R&D Systems) were used to determine the concentration of the respective proteins in the supernatant after appropriate dilution according to the manufacturers’ instructions. Plates were analyzed with a Spark 10M plate reader (TECAN).

### 2.7 Generation of *scamp3* knock-out zebrafish strains

*Scamp3* knock-out zebrafish lines were generated by CRISPR/Cas9-induced genome editing as described before ^27,28^. Briefly, the guide RNA TGGGTTGTAGAGGTCCAGCGTGG targeting exon 2 of the *scamp3* gene was injected into fertilized one cell stage AB wild-type embryos together with Cas9 protein (Integrated DNA Technology). Zebrafish of the F_0_ generation were raised and outcrossed with Tg(*fli1:gfp;lyz:dsred*) zebrafish. Offspring of the F_1_ generation were genotyped. DNA isolation and PCR for genotyping was performed with the PCRBIO Rapid Extract PCR Kit (PCR Biosystems) according to the manufacturer’s instructions using the primers 5’-ACATTCTCAGATCCACCA-3’ (forward) and 5’-CACTGTCAAAAAGTAGGGTT-3’ (reverse). PCR samples were sequenced to confirm the *scamp3* mutations. Two homozygous *scamp3* KO zebrafish were selected for further analyses. *Scamp3* KO 01 (Tg(*fli1:gfp;lyz:dsred*;*scamp3*^mde405^)) harbored a 39 base pair (bp) insertion, *scamp3* KO 02 (Tg(*fli1:gfp;lyz:dsred*;*scamp3*^mde406^)) a 7 bp deletion. The scamp3 KO 02 line was additionally outcrossed into the Casper background (Tg(*fli1:gfp;lyz:dsred*;*scamp3*^mde407^))^29^. All lines were bred to homozygosity.

Zebrafish larvae were kept in E3 medium (5 mM NaCl, 0.33 mM CaCl_2_, 0.33 mM MgSO_4_, 0.17 mM KCl, 0.00003% methylene blue) at 28.5 °C. 1-phenyl 2-thiourea (0.003%, Sigma-Aldrich) was added to the medium 24 h post fertilization to prevent pigmentation. Raising and housing of adult zebrafish and all described experiments were performed in accordance with animal protection standards of the Ludwig-Maximilians-Universität München and approved by the government of Upper Bavaria (Regierung von Oberbayern).

### 2.8 Granularity of zebrafish neutrophils

Neutrophil granularity was analyzed in neutrophils isolated from whole kidneys of adult *scamp3* WT and KO 01 zebrafish in the AB background. Additionally, *scamp3* KO 02 in the Casper background with appropriate WT controls were used. Adult zebrafish were euthanized by an overdose tricaine. The head was removed, a long ventral incision was made, and internal organs were carefully removed. The kidney was carefully peeled off the dorsal wall, transferred into 1 ml PBS containing 5% FCS, and immediately placed on ice. For the generation of a single cell suspension, the whole kidney marrow was gently homogenized through a 40 µm cell strainer using a rubber-coated syringe plunger and filtered into 10 ml ice-cold PBS. Samples were centrifuged at 200 x g for 5 min at 4 °C and resuspended in PBS containing SYTOX™ Red Dead Cell Stain (5 nM, Invitrogen). Samples were filtered once more through a 35 µm cell strainer and incubated on ice for 15 min. Samples were analyzed using a CytoFLEX S flow cytometer (Beckman Coulter) with subsequent data analysis with FlowJo^TM^ software (BD Biosciences). Granularity of DsRed-positive neutrophils was assessed by the sideward scatter.

### 2.9 Statistics

Data are represented as means±sd, unless stated otherwise, from three or more independent experiments. Statistical significance was determined by one-way or two-way ANOVA with Šídák’s or Tukey’s multiple comparisons test or Student’s t test using GraphPad Prism 10 (GraphPad Software). *P* values <0.05 were considered significant, *P* values are marked as * <0.05, ** <0.01, *** <0.001, and **** <0.0001.

## 3 Results

### 3.1 Host defense function of *Scamp3*-deficient Hoxb8 cell-derived neutrophils

To decipher the functional role of SCAMP3 for neutrophil biology, we generated *Scamp3* control (CTRL) and knockout (KO) Hoxb8 cells using CRISPR/Cas9 technique (Fig. 1A). KO 01 carries a point-mutation and KO 02 harbors a 32 bp deletion, both resulting in premature stop codons after 18 and 7 amino acids, respectively. Hoxb8 cells that underwent the same procedure but without mutations in *Scamp3* were used as CTRL cells. The absence of SCAMP3 in KO 01 and 02 was verified by Western blot technique. Correct differentiation of *Scamp3* CTRL and KO Hoxb8 cells into Hoxb8 cell-derived neutrophils (dHoxb8 cells) was analyzed by May-Grünwald-Giemsa staining and flow cytometry (Fig. 1B and C). *Scamp3* KO dHoxb8 cells showed the typical segmented nucleus similar to *Scamp3* CTRL dHoxb8 cells (Fig. 1B). Accordingly, *Scamp3* KO Hoxb8 cells downregulated the progenitor markers CD117 and CD34 and upregulated the neutrophil markers lymphocyte antigen 6G (LY6G) and C-X-C chemokine receptor type 2 (CXCR2) on the cell surface during differentiation similar to *Scamp3* CTRL dHoxb8 cells as expected (Fig. 1C). Additionally, no compensatory protein regulation of the other SCAMP protein members SCAMP1, 2, 4 or 5 was detected in *Scamp3* KO dHoxb8 cells (Supplementary Fig. 1A-G).

**Figure 1.**
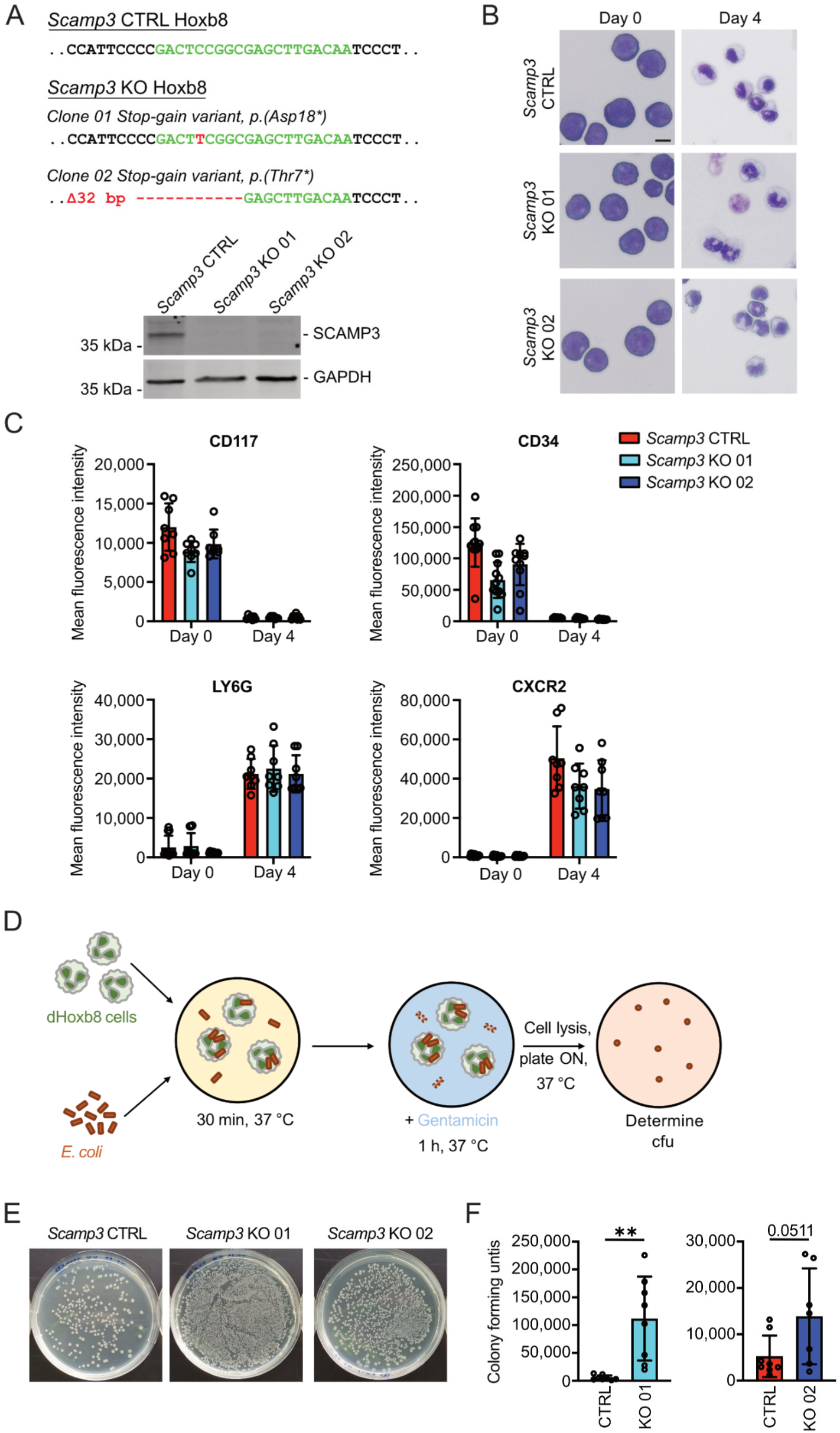
Characterization of *Scamp3* control (CTRL) and KO dHoxb8 cells. (A) Upper panel: Genomic sequence of *Scamp3* CTRL, KO 01 and KO 02 Hoxb8 cells generated by CRISPR/Cas9 technique. gRNA sequence, green. Deleted or mutated nucleotides, red. Lower panel: Representative image of a Western blot of whole-cell lysates of *Scamp3* CTRL, KO 01 and KO 02 dHoxb8 cells. SCAMP3 and GAPDH were detected by immunoblotting using specific antibodies. n=5. (B) May-Grünwald-Giemsa staining at day 0 and 4 of differentiation of indicated cell and genotypes. Scale bar, 10 µm. n=3. (C) Cell surface expression of CD117, CD34, LY6G and CXCR2 was analyzed by flow cytometry using specific fluorescently labeled antibodies. Mean fluorescence intensity is shown. n≥7. (D) Workflow of gentamycin assays to identify phagocytosed, surviving bacteria. (E-F) Representative images (E) and quantification (F) of colony forming units (cfu) in indicated genotypes. n≥7.

Next, we analyzed host defense functions of *Scamp3* KO dHoxb8 cells. To this end, bacterial killing capacity was studied by incubation of dHoxb8 cells with *E. coli* for 30 min to allow phagocytosis of bacteria and subsequent treatment with gentamycin for 60 min to kill the remaining extracellular, non-phagocytosed bacteria (Fig. 1D). Subsequently, dHoxb8 cells were lysed and plated onto agar plates for culture of phagocytosed but viable *E. coli.* A significantly higher number of colony forming units (cfu) was obtained from *Scamp3* KO 01 and 02 dHoxb8 cells compared to *Scamp3* CTRL dHoxb8 cells (Fig. 1E and F). At the same time, the number of phagocytosing cells positive for *E. coli* as well as the amount of phagocytosed bacteria did not differ between *Scamp3* CTRL, KO 01 and 02 dHoxb8 cells (Supplementary Fig. 2A and B). These data indicate an impaired bacterial killing capacity of dHoxb8 cells in the genetic absence of *Scamp3*.

### 3.2 Granule proteins and degranulation in *Scamp3* CTRL and KO dHoxb8 cells

To decipher the underlying mechanism of impaired host defense in *Scamp3*-deficient dHoxb8 cells, the proteome of *Scamp3* CTRL, KO 01 and 02 dHoxb8 cells was analyzed by mass spectrometry (MS). Almost 4000 proteins per sample were quantified (Supplementary Fig. 3A). The principal component (PC) 1 analysis revealed a clear separation of Hoxb8 (day 0) and dHoxb8 (day 4) cells due to maturation into mature neutrophils as expected (Supplementary Fig. 3B). Furthermore, a separation of *Scamp3* CTRL to *Scamp3* KO 01 and 02 dHoxb8 cells was visible in PC 2, demonstrating a clear change of the proteome composition in the genetic absence of *Scamp3*. Gene ontology enrichment analyses of biological processes and cellular compartments revealed reduced regulation of vesicle mediated transport, secretory vesicles, secretory granules, endocytic vesicles, and immune effector processes, amongst others, indicating a possible effect on vesicles and neutrophil granules by SCAMP3 deficiency (Fig. 2A). Indeed, granularities of *Scamp3* KO 01 and 02 dHoxb8 cells were significantly reduced compared to CTRL conditions as evaluated via flow cytometric side scatter (SSC) (Fig. 2B). Strikingly, MS analyses revealed that protein amounts of the marker proteins myeloperoxidase (MPO), neutrophil elastase (NE) and cathepsin D for primary granules, lactoferrin (LTF), neutrophil gelatinase-associated lipocalin (NGAL) and olfactomedin-4 (OLFM4) for secondary granules, and neutrophil collagenase (MMP8), matrix metalloproteinase-9 (MMP9) and lysozyme C-2 (LYZ2) for tertiary granules were strongly downregulated in the genetic absence of *Scamp3* compared to CTRL conditions (Fig. 2C-E).

**Figure 2.**
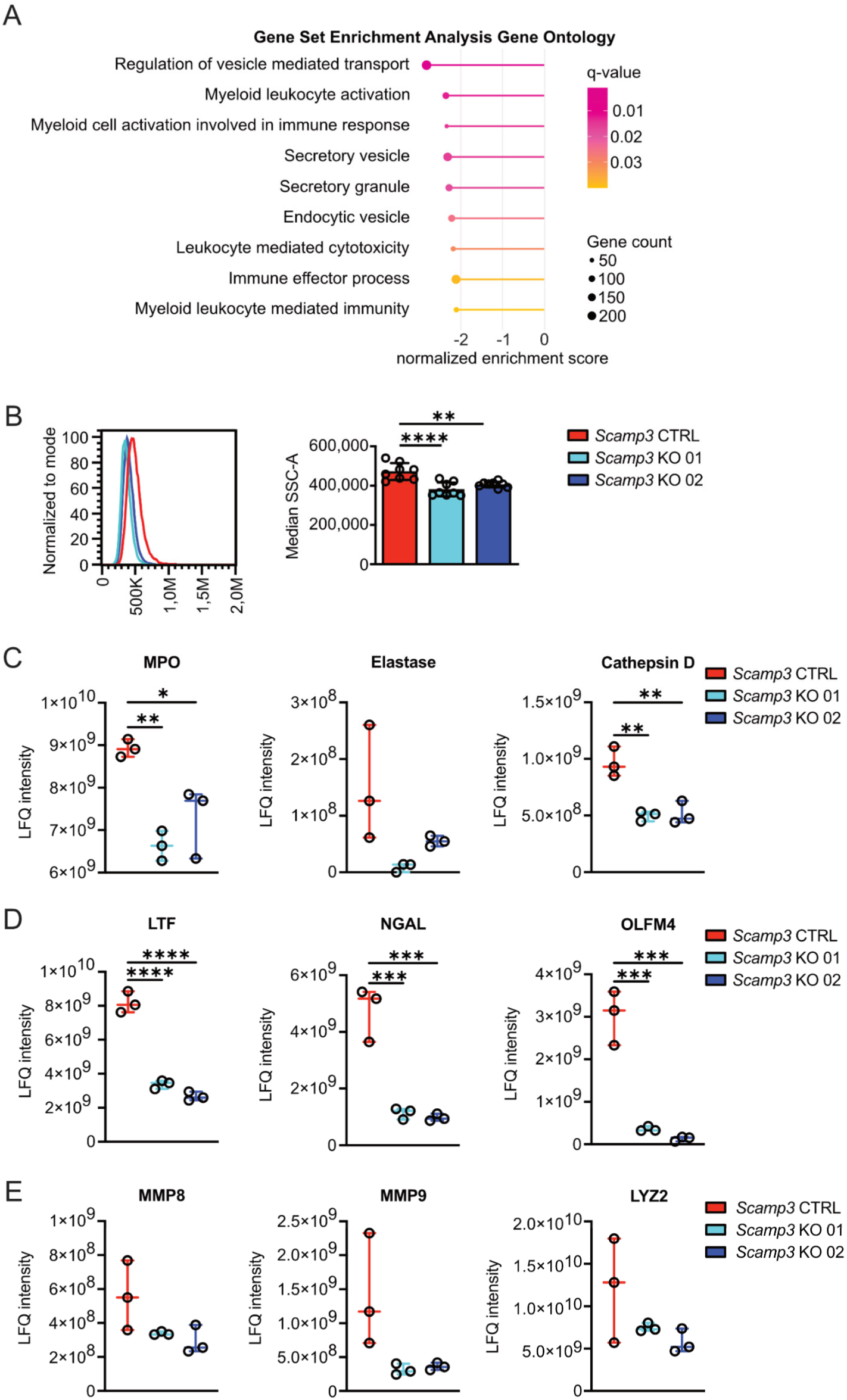
Granularity changes in *Scamp3*-deficient dHoxb8 cells. (A) Gene set enrichment analysis with gene ontology cellular components and biological processes of *Scamp3* KO 02 compared to *Scamp3* CTRL dHoxb8 cells from mass spectrometry data. (B) Flow cytometry histograms (left panel) and quantitative analysis (right panel) of SSC-A as a measure for granularity of dHoxb8 cells of indicated genotypes. n≥4. (C-E) Expression levels of marker proteins MPO, elastase and cathepsin D for primary granules (C), LTF, NGAL and OLFM4 for secondary granules (D) and MMP8, MMP9 and LYZ2 for tertiary granules (E) analyzed by mass spectrometry. n=3.

Downregulation of NE as specific marker for primary granules and MMP9 as specific marker for tertiary granules was not as pronounced as for the secondary granule markers. Therefore, reduced expression of NE and MMP9 were verified by Western blot technique and clearly, protein levels of NE and MMP9 were significantly downregulated in *Scamp3* KO dHoxb8 cells compared to *Scamp3* CTRL dHoxb8 cells (Fig. 3A and B). Next, we investigated whether the reduced protein amount affected degranulation. Indeed, all three marker proteins, MPO, LTF and MMP9, were reduced in the supernatant upon stimulation with fMLP and cytochalasin D (Fig. 3C-E), indicating a diminished degranulation of granule proteins in the genetic absence of *Scamp3.* Similarly, several membrane proteins of granules including CD63, carcinoembryonic antigen-related cell adhesion molecule 1 (CEACAM-1) and sialic acid-binding Ig-like lectin 5 (SIGLEC-5) in primary, secondary and tertiary granules, respectively were also downregulated (data not shown).

**Figure 3.**
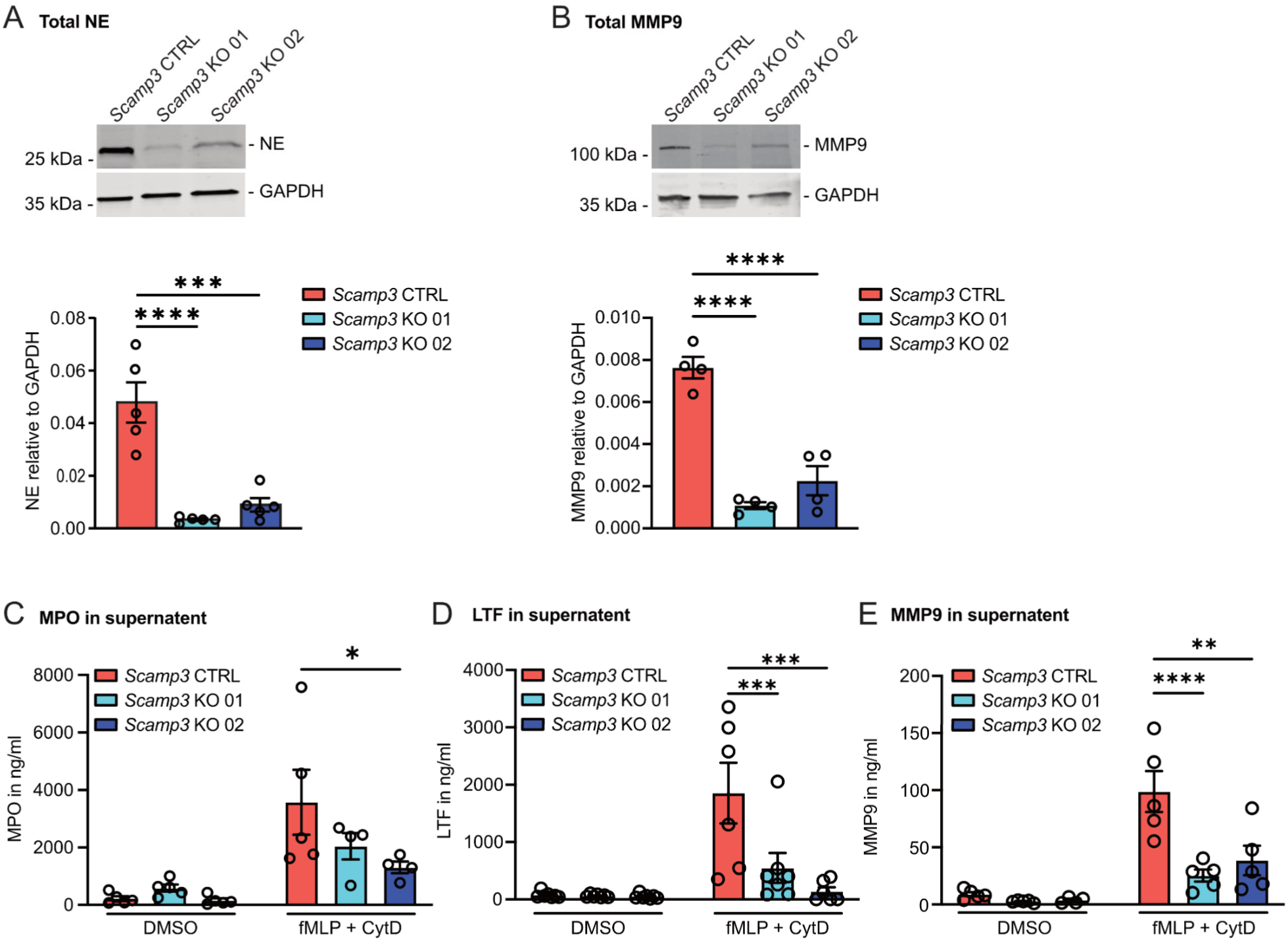
Granule proteins and degranulation in *Scamp3* dHoxb8 cells. (A, B) Representative Western blots (upper panel) and quantification (lower panel) of total protein amount of NE (A) and MMP9 (B) in whole cell lysates from *Scamp3* CTRL, KO 01 and 02 dHoxb8 cells. Indicated proteins were detected with specific antibodies. GAPDH was used as loading control. n≥4. Mean ± SEM. (C, D, E) Degranulated MPO (C), LTF (D) and MMP9 (E) upon stimulation with fMLP and cytochalasin D (CytD) analyzed with an ELISA. n≥4. Mean ± SEM.

To analyze a possible effect of the loss of SCAMP3 on secretory vesicles, surface levels of the β_2_ integrin subunits CD11b and CD18 were studied under steady state conditions and after fMLP stimulation which causes fusion of secretory vesicles with the plasma membrane and accordingly upregulation of CD11b and CD18 on the cell surface (Supplementary Fig. 4A, B).^4,30,31^ However, no significant difference of upregulated CD11b on the cell surface was found between *Scamp3* CTRL, KO 01 and KO 02 dHoxb8 cells. The MS data confirmed similar total protein levels of CD11b and CD18 in these cells (Supplementary Fig. 4C). Furthermore, induction of neutrophil adhesion as well as mechanotactic migration under flow conditions that depend on rapid release of secretory vesicles to allow β_2_ integrin-dependent neutrophil adhesion and migration were not affected by the loss of SCAMP3, indicating that SCAMP3 is dispensable for proper generation and release of secretory vesicles (Supplementary Fig. 4D, E).^4^

### 3.3 Host defense function of *scamp3* WT and KO zebrafish larvae

The role of SCAMP3 for host defense *in vivo* was studied in *scamp3* WT and KO zebrafish larvae. We performed sequence alignments and found that human, murine and zebrafish SCAMP3 protein has a very similar structure with a long N-terminus, a proline rich region and four transmembrane domains (Supplementary Fig. 5A). As expected, the sequence identity and similarity between human SCAMP3 and murine SCAMP3 was very high with 90.26% and 92.84%, respectively (Supplementary Fig. 5B). Zebrafish Scamp3 shared 59.38% identity and 70.74% similarity with human SCAMP3. Strikingly, zebrafish Scamp3 also harbored the conserved NPF, PPxY, PSAP and PTEP protein motifs indicating that SCAMP3 might share similar functions between human, mice and zebrafish and that the zebrafish is a suitable model to study SCAMP3 (Supplementary Fig. 5C).

Next, we generated *scamp3* KO zebrafish using CRISPR/Cas9 technique by targeting exon 2 of the *scamp3* gene in the transgenic zebrafish line *Tg(fli:gfp;lyz:dsRed)* expressing green fluorescent protein (GFP) under the promoter *fli1* and DsRed under the promoter *lyz* to specifically label endothelial cells and neutrophils, respectively (Fig. 4A).^27,32–34^ *Scamp3* KO 01 and *scamp3* KO 02 were chosen for further analyses. *Scamp3* KO 01 harbored a 38 bp insertion and *scamp3* KO 02 a 7 bp deletion, both resulting in premature stop codons after 38 and 44 bp, respectively. Both *scamp3* KO zebrafish lines were viable and fertile. The average total neutrophil counts of 126±33 neutrophils per larva in 3 dpf *scamp3* WT zebrafish larvae were not different to those of *scamp3* KO 01 with 128±19 and *scamp3* KO 02 with 114±26 neutrophils per larva (Fig. 4B).

**Figure 4.**
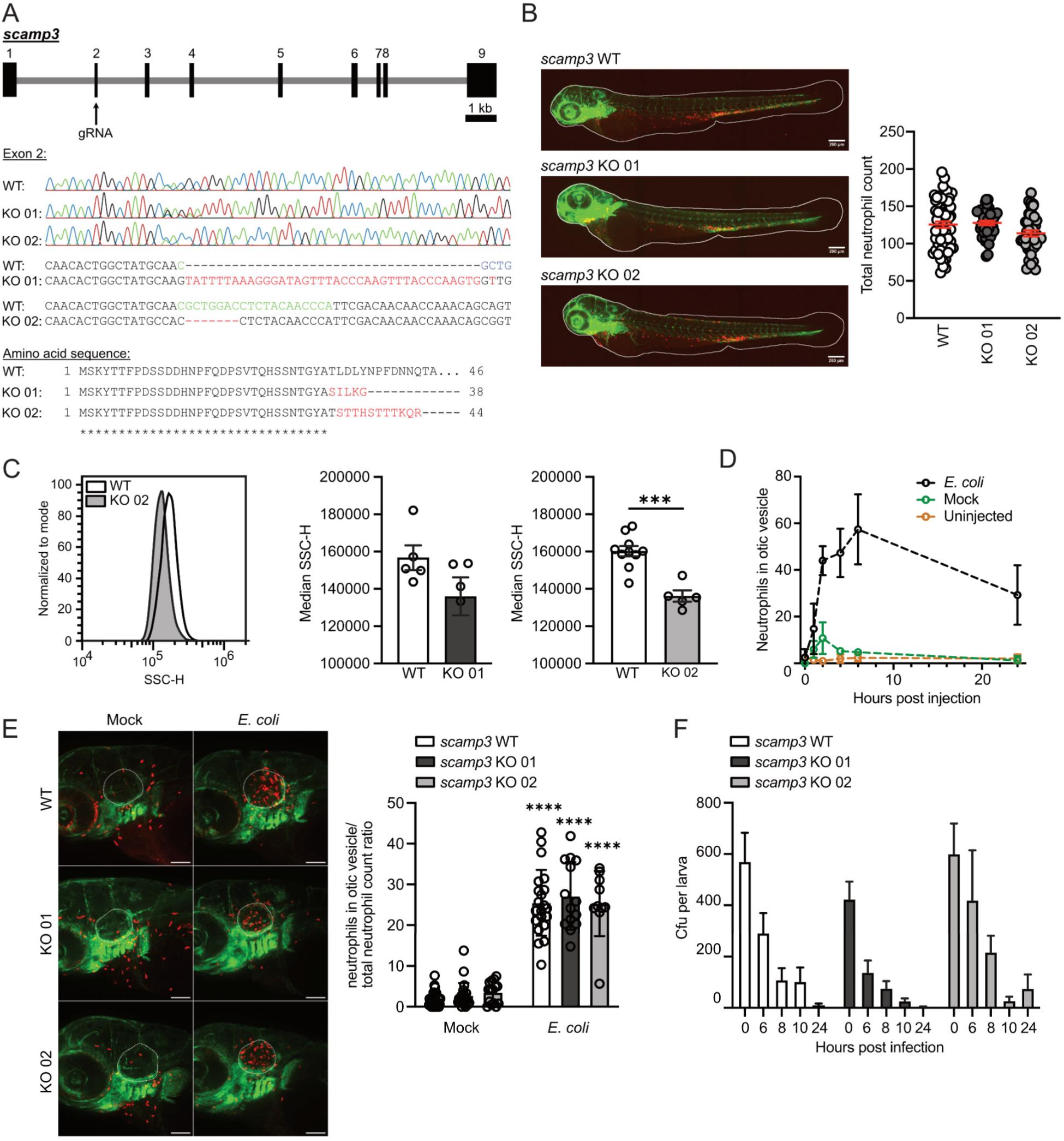
Characterization of neutrophils in *scamp3* WT and KO zebrafish larvae. (A) Generation of *scamp3* KO zebrafish using CRISPR/Cas9. Upper panel: Schematic of the zebrafish *scamp3* genomic organization with gRNA1 exon 2 target site. Middle panel: Sequencing traces of *scamp3* WT, KO 01 and 02. gRNA sequence, green. Nucleotides mutated or deleted around mutation site, red. Lower panel: Predicted amino acid sequence of *scamp3* KOs aligned to the first 46 amino acids of *scamp3* WT sequence. Identical (*) and altered (red) amino acids are indicated. Predicted protein length in *scamp3* KO 01 is 38 amino acids, in *scamp3* KO 02 44 amino acids. (B) Representative microscopic images of zebrafish larvae of indicated genotypes 3 dpf (left panel) and quantification of total neutrophils (right panel). Scale bar, 200 µm. n≥40. Mean ± SEM. (C) Flow cytometry histograms (left panel) and quantitative analysis (right panel) of SSC-H as a measure for granularity of neutrophils isolated from adult zebrafish kidney of indicated genotypes. n≥5. (D) Neutrophils recruited to the otic vesicle 0, 1, 2, 4, 6 and 24 h post-infection with 2,000 cfu *E. coli*. n=2 (0 h), n≥2 (1 h), n≥3 (2 h), n≥3 (4 h), n≥3 (6 h), n=4 (24 h). Mean ± SEM. (E) Representative microscopic images (left panel) and quantification (right panel) of neutrophils recruited to the otic vesicle 4 h post infection in zebrafish larvae 3 dpf of indicated genotypes. n≥10. Scale bars, 100 µm (F) Cfu per larva of *scamp3* WT, KO 01 and KO 02 zebrafish larvae 3 dpf infected with 2,000 cfu *E. coli* at the otic vesicle. n≥12. Mean ± SEM.

To verify our *in vitro* findings, the granularity of neutrophils was analyzed *ex vivo*. For this, whole kidney as the site of hematopoiesis in adult zebrafish was isolated, dissected into a single cell solution and analyzed via flow cytometry. As described elsewhere, myelomonocytes were identified using forward scatter (FSC) and SSC.^35^ From this population neutrophils were identified as DsRed-positive cells. The granularity of neutrophils was assessed by their median SSC (Fig. 4C). Here, *scamp3* KO 01 and *scamp3* KO 02 neutrophils were characterized by a reduced granularity compared to *scamp3* WT neutrophils, confirming our findings in dHoxb8 cells.

In order to determine whether the reduced neutrophil granularity dampens bacterial killing *in vivo*, we first investigated whether *scamp3*-deficient neutrophils were able to migrate to sites of infection. To determine the appropriate time-point for these experiments, a time-course of neutrophil recruitment to the otic vesicle of 3 dpf larvae infected with approximately 2,000 cfu *E. coli* was generated (Fig. 4D). Injections with PBS (mock) were used as control reflecting the trauma-induced recruitment of neutrophils. Zebrafish larvae that were left untreated served as control displaying the number of neutrophils in the otic vesicle at steady state. Over the time course of 24 h, a maximum of 3±1 neutrophils were in the otic vesicle in untreated zebrafish larvae (Fig. 4D). Mock injections led to an average increase to 6±4 recruited neutrophils at 1 h post injection (hpi) with a peak of 11±7 recruited neutrophils at 2 hpi. Subsequently, the number of neutrophils at the otic vesicle went back to control levels. In contrast, injection of *E. coli* led to the recruitment of 15±11 neutrophils 1 hpi, 44±6 neutrophils 2 hpi, 47±10 neutrophils 4 hpi and peaked with 57±15 neutrophils 6 hpi. After 24 h the number of neutrophils decreased to 29±13 at the otic vesicle. For further experiments, the time-point of 4 hpi was chosen where the recruitment had not reached a plateau yet.

Three dpf *Scamp3* WT and KO zebrafish larvae were analyzed 4 hpi and for accuracy the number of recruited neutrophils is given relative to the respective total neutrophil count of each individual larva. Compared to mock-injected larvae, the number of recruited neutrophils to *E. coli*-infected otic vesicles increased significantly from 2±2% to 25±8% in *scamp3* WT larvae, from 3±3% to 27±8% in *scamp3* KO 01 larvae and from 4±3% to 25±8% in *scamp3* KO 02 larvae (Fig. 4E). Further, no differences in the recruited neutrophils between *scamp3* WT and KO zebrafish larvae were observed indicating that *scamp3*-deficient neutrophils are able to migrate to sites of infection and that Scamp3 is dispensable for neutrophil migration during acute, infectious inflammation in zebrafish larvae. Next, the potential effect of reduced neutrophil granularity on bacterial killing was evaluated by determining the cfu of injected *E. coli* at 6, 8, 10 and 24 hpi (Fig. 4F). However, 3 dpf *scamp3* KO 01 and KO 02 zebrafish larvae were able to combat the infection similar to *scamp3* WT zebrafish larvae. These data indicate that the remaining defense mechanisms seem to be still intact and sufficient to kill invading bacteria and that they are regulated independently of SCAMP3. Alternatively, reduced granularity of neutrophils may be compensated by other components of the innate immune system *in vivo*.

In summary, we found that the genetic absence of *Scamp3* in dHoxb8 cells resulted in impaired pathogen killing due to reduced granularity and granule proteins. Further, *scamp3* KO zebrafish also showed neutrophils with a reduced granularity but were still able to combat invading *E. coli* indicating that SCAMP3 might be critical for granule homoeostasis and bacterial killing *in vitro* but the loss of SCAMP3 can be compensated *in vivo*.

## 4 Discussion

In the present study, we identified SCAMP3 as a novel regulator of granule homoeostasis in neutrophils. Bacterial killing assays revealed an impaired killing ability in the genetic absence of *Scamp3 in vitro*. Moreover, MS analyses revealed a strong reduction in granule proteins. Further analyses verified these data and unraveled that degranulation of primary, secondary and tertiary granules was also diminished in the absence of *Scamp3*. Interestingly, secretory vesicles seemed not to be altered in the absence of SCAMP3 as indicated by similar total protein amounts of β_2_ integrin subunits CD11b and CD18 and cell surface levels of CD11b and CD18 in unstimulated *Scamp3* CTRL and KO dHoxb8 cells. CD11b and CD18 are located in secretory vesicles, tertiary and secondary granules and stimulation with fMLP induces fusion of secretory vesicles with the plasma membrane.^30,36–38^ In accordance with this, CD11b cell surface levels were not altered in the absence of SCAMP3 upon fMLP stimulation. Notably, interaction of neutrophils via selectin ligands such as P-selectin glycoprotein ligand 1 (PSGL-1) with the inflamed endothelium induces neutrophil rolling and the release of secretory vesicles and with this the upregulation of β_2_ integrins and chemokine receptor CXCR2 on the cell surface leading to neutrophil adhesion.^31^ In line with this, β_2_ integrin-dependent recruitment steps including induction of adhesion and mechanotactic migration were not altered in *Scamp3*-deficient dHoxb8 cells compared to *Scamp3* CTRL dHoxb8 cells. In the present study, the experiments were carried out in a reductionist approach with flow chambers coated with P-selectin, intercellular adhesion molecule 1 (ICAM-1) and C-X-C motif chemokine ligand 1 (CXCL-1) resembling the inflamed endothelium. Thus, our data suggest that SCAMP3 is indispensable for granule formation and degranulation but dispensable for secretory vesicle homeostasis and function.

The regulation of granules but not secretory vesicles by SCAMP3 may be due to their different formation pathways. Secretory vesicles are from an endocytic origin.^36^ In contrast, granules originate from the biosynthetic pathway.^39^ During this process, protein cargo is routed from the *trans* Golgi network (TGN) to granules upon translation at the ER and post-translational modification in the Golgi complex. This trafficking is in part mediated by adaptor protein complexes (APs) 1, 3 and 4. Here, AP-3 directs cargo to granules either directly from the TGN or from sorting endosomes. SCAMP3 is localized in almost all membranes of the cell including the plasma membrane and endolysosomal compartments but is especially accumulated in the membranes of the TGN.^11^ Here, SCAMP3 strongly colocalizes with AP-3 in in the rat kidney cell line NRK. In the same study, the authors speculate that SCAMP3 functions in organizing membrane budding sites. In addition, SCAMP3 may be critical for directing proteins into granules at the TGN in neutrophils. This is in line with findings by Norris et al. who found a reduced content of insulin in TGN-derived insulin secretory granules in murine *Scamp3* knockdown β-cells.^40^ With regard to the observed reduced degranulation in our study, SCAMP3 has been shown to also regulate exocytosis in endocrine PC12 cells indicating that reduced degranulation in the absence of SCAMP3 might not only be a result of reduced granule proteins but also potentially due to impaired exocytosis of granules.^41^

Granule protein deficiencies can result in impaired host defense.^38,42^ A variety of diseases has been described where gene mutations affect granule proteins and hence pathogen clearance. Chronic granulomatous disease (CGD) is an immunodeficiency induced by mutations affecting the nicotinamide adenine dinucleotide phosphate (NADPH) oxidase complex or its assembly leading to the inability of neutrophils, monocytes and macrophages to produce reactive oxygen species or superoxide anions.^43^ The two subunits gp91*^phox^* and p22*^phox^* are located in granule membranes and mutations, amongst others in *CYBB*, the gene encoding for gp91*^phox^*, or *CYBA*, encoding for p22*^phox^*, result in CGD.^4,44^ Patients with CGD suffer from recurrent specific bacterial and fungal infections. Leukocyte adhesion deficiency type I is induced by mutations in *ITGB2*, the gene encoding for CD18, the β-subunit of β_2_ integrins which is present in secretory vesicles, tertiary and secondary granules.^45^ The genetic absence of CD18 results in the inability of leukocytes to firmly adhere and migrate on the inflamed endothelium and hence to reach the site of inflammation leading to neutrophilia and life-threatening bacterial and fungal infections. Specific granule deficiency (SGD) is a disease that is caused by defects in the transcription factor CCAAT/enhancer binding protein (C/EBP) ε, resulting in a lack of specific granules in neutrophils.^4,46,47^ A hallmark of SGD are the deficiency or very low levels of lactoferrin. Patients with SGD are highly susceptible to bacterial infections of the skin and mucous membranes, and neutrophils from these patients show an impaired bactericidal activity and defective chemotaxis. In line with the described immunodeficiencies, *Scamp3*-deficient dHoxb8 cells show reduced bactericidal activities *in vitro*. Although C/EBPε expression levels were slightly reduced in the MS analysis (data not shown), the effect of reduced granule proteins is not restricted to secondary but rather to all granules pointing towards a more general role of SCAMP3 in granule protein production and distribution.

Unexpectedly, we were not able to verify our data *in vivo* as bacterial killing capacities of *E. coli*-infected *scamp3* KO zebrafish larvae were comparable to those of *scamp3* WT zebrafish larvae. However, similar findings have been reported for MPO deficiencies in humans. MPO deficiencies in humans do not necessarily cause disease and MPO-deficient humans revealed by screening analyses appear healthy.^42,48–50^ Only in some instances MPO-deficient patients, often those with other co-morbidities including diabetes, present with symptoms such as disseminated *Candida* infection. Interestingly, *in vitro* studies analyzing the microbicidal capabilities of MPO-deficient neutrophils revealed a significantly reduced killing capacity of *Staphylococcus aureus* and *Candida albicans* by MPO-deficient neutrophils while killing of *E. coli* was not impaired.^51–55^ However, MPO-deficient mice have an increased susceptibility to a range of pathogens including *Candida* and *Klebsiella*.^48,56–58^ Studies in MPO-deficient mice also revealed that the loss of MPO can be compensated for low but not for high infectious doses of pathogenic fungi.^59,60^ These data indicated that neutrophils or hosts deficient for one defense component were still able to combat pathogens in general, but were susceptible to only a specific selection of pathogens. With regard to our findings, where no considerable difference of bacterial clearance between *scamp3* WT and KO zebrafish larvae upon injection of *E. coli* was observed, it is tempting to speculate that the defense against this bacterial species was still intact or the bacterial burden was still low enough. Apart from that, other cells of the innate immune system such as macrophages might have taken over the task of bacterial elimination.

In conclusion, our data indicate that SCAMP3 may function in proper formation of granules and degranulation. The reduction in granularity and degranulation resulted in impaired bacterial killing of *E. coli in vitro*. Interestingly, *scamp3*-deficient zebrafish larvae did not show a bacterial killing defect indicating that SCAMP3 is specifically required for neutrophil granule formation and function whereas other defense mechanism seem to be independent of SCAMP3. These residual defense components were sufficient to eliminate the here applied *E. coli* strain. Recently, SCAMP3 has emerged as a viral target for successful replication.^17,18^ As a result, SCAMP3 inhibitors for treatment of chronic hepatitis b virus (HBV) infection are currently under investigation for destabilization HBV cccDNA which is necessary for its replication as well as to reduce SCAMP3 mRNA levels.^61^ In this context, the effect of SCAMP3 inhibition on neutrophil defense function has to be carefully considered.

## Supporting information

BaderEtAl_SupplementaryMaterial

## Acknowledgments

The authors thank Jennifer Truong, Tanja Vlaovic, Tanja Weißer, Ulrike Wilhelm-Forster, Enrique de Vega Gómez, and Sabine Schlink for excellent technical assistance. The authors are grateful to Dr. Steffen Dietzel (Core Facility Bioimaging, Biomedical Center, LMU Munich) for the support with fluorescence microscopy. We acknowledge the Core Facility Flow Cytometry at the Biomedical Center, LMU Munich, for providing equipment and expertise.

## 6 Author contributions

AB carried out research, analyzed data and wrote the manuscript. JG and NR carried out research and analyzed data. AZ generated *Scamp3* CTRL and KO Hoxb8 cells and analyzed data. DS and DRE analyzed mass spectrometry data. BS carried out research and analyzed data. IF performed mass spectrometry and analyzed data. BW designed the research and analyzed data. DM-B designed the research, carried out research, analyzed data and wrote the manuscript.

