## Supplementary material for "SCAMP3 is essential for proper formation and function of neutrophil granules": BaderEtAl_SupplementaryMaterial

**[3]German Center for Neurodegenerative Diseases (DZNE), Munich, Germany.**

**[4]Protein Analysis Unit, Biomedical Center, LMU Munich, 82152 Planegg-Martinsried, Germany**

**# Correspondence:**

**Daniela Maier-Begandt**

Institute of Cardiovascular Physiology and Pathophysiology
Biomedical Center
LMU Munich
Großhadernerstr. 9
82152 Planegg-Martinsried
Germany

### Supplementary Methods

#### 1.1 May Grünwald Giemsa staining

Cytospins were prepared from 5 x $10^4$ Hoxb8 cells per slide at day 0 or day 4 of differentiation using a Cellspin III Cytocentrifuge (THARMAC). Cells were stained first with May-Grünwald solution (Carl Roth) for 3 min and then with 1.3% Giemsa solution (Merck Millipore) for 20 min. Imaging was performed with a bright field Leica DM2500 microscope equipped with a DMC2900 CMOS camera and a 100 x/1.40 oil immersion objective (Leica).

#### 1.2 Immunoblotting

Differentiated Hoxb8 cells were lysed using M-PER™ Mammalian Protein Extraction Reagent (Thermo Fisher Scientific) containing 1 mM diisopropyl fluorophosphate, 100 mM sodium orthovanadate, 100 mM sodium fluoride, and protease inhibitor cocktail (Sigma-Aldrich). The total protein concentration was estimated using a NanoDrop™ 2000 microvolume spectrophotometer at 280 nm (Thermo Fisher Scientific). 10-30 µg total protein were separated in 4-20% Mini-PROTEAN® TGX Stain-free™ protein gels (Bio-Rad) before transfer onto nitrocellulose membranes. Blocking was performed with 5% non-fat dried milk powder in TBS containing 0.1% Tween-20 (TBS-T) or 50% Intercept® blocking buffer TBS (LI-COR) in TBS-T. Primary antibodies targeting SCAMP3 (1:1000, PA5-21428, Invitrogen), neutrophil elastase (1:1000, MAB4517, R&D Systems), MMP9 (1:1000, MA5-15886, Invitrogen), SCAMP1 (1:500, PA1-739, Invitrogen), SCAMP2 (1:500, PA1-734, Invitrogen), SCAMP4 (1:500, HPA043284, Atlas Antibodies), SCAMP5 (1:1000, PA1-737, Invitrogen), and GAPDH (1:5000, Merck Millipore) were added over night in blocking buffer. Secondary IRDye 800CW donkey anti-rabbit (1:10000, 926-32213, LI-COR Biosciences) and IRDye 680RD donkey anti-mouse antibodies (1:10000, 926-68072, LI-COR Biosciences) were incubated for 1 h at room temperature in blocking buffer. Detection was performed with an Odyssey® CLx Imaging system (LI-COR Biosciences) equipped with Image Studio software (LI-COR Biosciences).

#### 1.3 Bacterial uptake assay

For assessment of bacterial uptake, 2 x $10^5$ differentiated Hoxb8 cells were incubated with *E. coli* MG1655 pEB2-E2-Crimson with an MOI of 10 for 30 min at 37 °C in PBS containing 10% FCS. Where indicated cells were pre-treated with 10 µg/ml cytochalasin D for 15 min at 37 °C. Afterwards, cells were centrifuged, resuspended in LIVE/DEAD™ Fixable Violet Dead Cell Stain (Invitrogen) in PBS, and incubated on ice for 15 min before fixation with 2% formaldehyde in PBS for 15 min on ice. Samples were analyzed using a CytoFLEX S flow cytometer (Beckman Coulter) with subsequent data analysis with FlowJo™ software (BD Biosciences).

#### 1.4 Flow chamber assays

Induction of adhesion of dHoxb8 cells was analyzed in µ slides VI$^{0.1}$ (Ibidi) coated with recombinant murine (rm) P-selectin (5 µg/ml), rm intercellular adhesion molecule 1 (ICAM-1, 3 µg/ml), and rm C-X-C motif chemokine ligand 1 (CXCL-1, 5 µg/ml) as described before.[1] 7.5 x $10^5$ dHoxb8 cells were perfused into the flow channels in adhesion medium with a constant flow of 1 dyne/cm² for a duration of 9 min. Time-lapse images were acquired of 18 different fields of view using an Axiovert 200M microscope equipped with a Plan-Apochromat 20 x/0.75 NA objective, AxioCam HR digital camera, and a temperature-controlled environmental chamber (Zeiss). The number of adherent cells after 1, 3, 5, 7, and 9 min was counted manually using FIJI.

For the analysis of mechanotactic migration µ slides VI$^{0.1}$ (Ibidi) were coated with rmICAM-1 (3 µg/ml) and rmCXCL-1 (5 µg/ml). dHoxb8 cells were filled into the channels in adhesion medium and allowed to adhere for 5 min before application of a constant flow of 1 dyne/cm² for 10 min. Images were acquired every 5 s with the microscope described above. Migration tracks of individual cells were analyzed manually using FIJI with the Manual Tracking plugin (Fabrice P. Cordelières, Institute Curie) and the Chemotaxis Tool (Ibidi).

#### 1.5 Alignments

The protein sequences of human (*Homo sapiens*), mouse (*Mus musculus*), and zebrafish (*Danio rerio*) SCAMP3 were aligned with Clustal Omega[2], using the longest sequence from the UniProt database.[3] Identity and similarity were analyzed with the Sequence Manipulation Suite: Ident and Sim[4], using default settings.

#### 1.6 Microscopy of zebrafish larvae

For microscopy zebrafish larvae were euthanized by an overdose tricaine (0.3 mg/ml, Pharmaq Ltd) and fixed in 4% formaldehyde in PBS over night at 4 °C. After washing, larvae were mounted in 1.5% low-melting agarose and imaged with an upright spinning disc confocal laser microscope (Examiner, Zeiss) equipped with a confocal scanner unit CSU-X1 (Yokogawa Electric Corporation), a CCD camera (Evolve, Photometrics) and a 5x/0.15NA objective (N-Achroplan, Zeiss) and Slidebook 6.0.13 Software (3i). Total neutrophil numbers within the whole larvae and in the otic vesicle after bacteria injection were counted manually in a blinded manner using the Cell Counter plugin in FIJI[5].

#### 1.7 Bacteria injection into the otic vesicle of zebrafish larvae

Injection of bacteria into the otic vesicle of zebrafish larvae at 3 days post fertilization (dpf) was done as described before[6]. *E. coli* MG1655 carrying the plasmid pEB2-E2-Crimson were cultured as described above, centrifuged at 2000 x g for 5 min, and resuspended at a concentration of $10^9$ bacteria/ml in PBS containing 0.1% phenol red (Sigma-Aldrich).

Microinjection needles were prepared using glass capillaries (World Precision Instruments) which were pulled with a micropipette puller (Sutter Instrument) set to the following parameters: heat 450, pull 30, velocity 100, time 60, pressure 500, ramp 465. 3 µl of bacteria suspension was loaded into the injection needle and fastened onto a micromanipulator (World Precision Instruments) connected to a FemtoJet® 4x (Eppendorf). The tip of the needle was clipped using thin forceps and the correct volume per drop was adjusted by measuring the drop size with a micrometer scale (Edmund Optics). 2 nl of bacteria suspension containing approximately 2000 *E. col*i or phenol red solution only (mock) were injected into the otic vesicle of zebrafish larvae that were anesthetized with tricaine (0.08 mg/ml) in E3 medium. For quantification of neutrophil numbers at the otic vesicle immediately after injection, zebrafish larvae were mounted in 1.5% low-melting agarose in E3 medium containing 0.08 mg/ml tricaine and imaged as described above. Afterwards, zebrafish larvae were carefully placed into single wells of a 96 multiwell plate, washed twice in E3 medium, and incubated at 28.5 °C for indicated time points before euthanasia, fixation and imaging as described above. Survival of zebrafish larvae was observed up to 48 h after bacteria injection.

### 1.8 Bacterial killing in zebrafish larvae

Approximately 2000 *E. coli* MG1655 pEB2-E2-Crimson per larvae were injected into the otic vesicle of *scamp3* WT, KO 01, and KO 02 zebrafish larvae that were anesthetized with tricaine (0.08 mg/ml) in E3 medium. After injection, zebrafish larvae were carefully placed into individual wells of a 96 multiwell plate, washed twice in E3 medium, and incubated for 0, 6, 8, 10, and 24 h at 28.5 °C. Afterwards, larvae were euthanized, washed twice with PBS, and resuspended in 500 µl PBS containing 0.2% TritonTM X-100 (AppliChem). Larvae were homogenized using a Pellet Pestle® Cordless Motor (Sigma-Aldrich) for 45 s. 50 µl of homogenates were plated onto LB agar plates containing 50 µg/ml kanamycin and incubated over night at 37 °C. *E. coli* colonies were counted manually.

**Supplementary Figure 1**

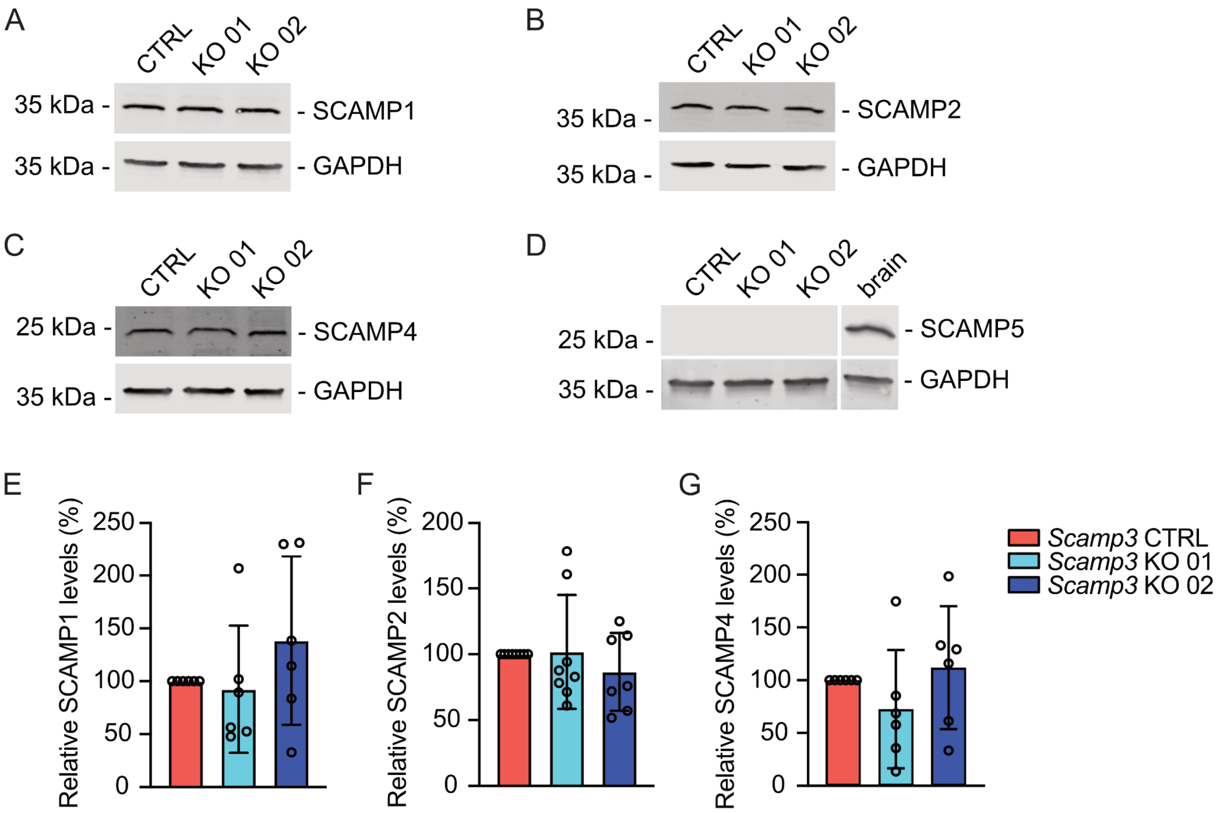


**Supplementary Figure 1.** Characterization of *Scamp3* control (CTRL) and KO dHoxb8 cells. (A-D) Representative images of Western blots of whole-cell lysates of *Scamp3* CTRL, KO 01 and KO 02 dHoxb8 cells. SCAMP1 (A), SCAMP2 (B), SCAMP4 (C), SCAMP5 (D) and GAPDH were detected by immunoblotting using specific antibodies. n≥5. (E-G) Expression levels of SCAMP1 (E), SCAMP2 (F) and SCAMP4 (G) quantified from Western blots from (A-C). n≥6.

**Supplementary Figure 2**

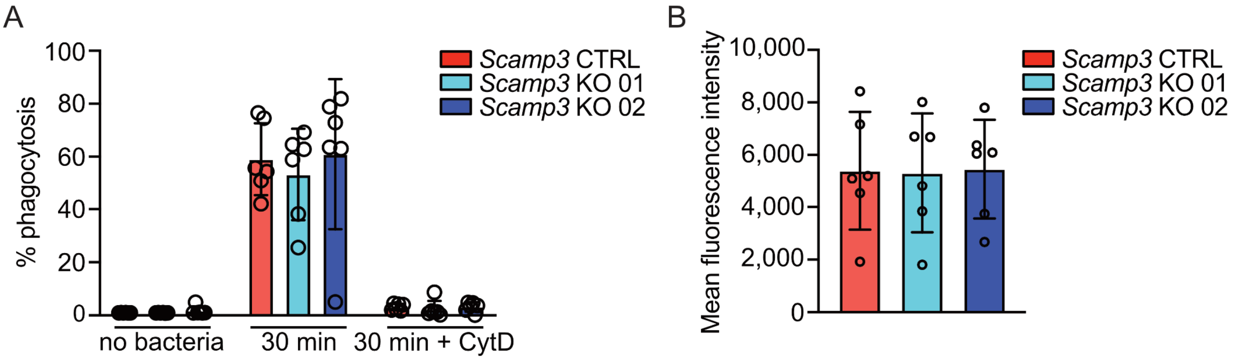


**Supplementary Figure 2.** Phagocytosis of live *E. coli* of *Scamp3* CTRL, KO 01 and KO 02 dHoxb8 cells. (A) % of phagocytosing cells (live cells, 100%) after 30 min incubation with *E. coli* and without bacteria as well as with *E. coli* in presence of cytochalasin D as control. n=6. (B) The amount of phagocytosed bacteria was evaluated by mean fluorescence intensity of E2 crimson-positive cells, fluorescence intensity values of cytochalasin D-treated control cells were subtracted. n=6.

**Supplementary Figure 3**

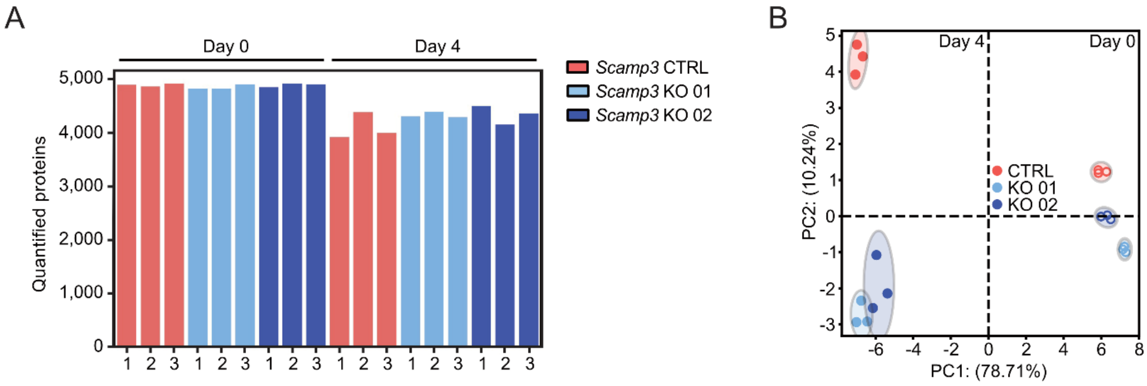


**Supplementary Figure 3.** Proteomic changes of Hoxb8 cells in the absence of SCAMP3. Cell lysates of *Scamp3* CTRL, KO 01 and KO 02 dHoxb8 cells were subjected to LC-MS/MS analysis. (A-B) Number of quantified proteins (A) and principal component analysis using proteins with a mutual information > 1.2 in *Scamp3* CTRL, KO 01 and 02 Hoxb8 cells on days 0 (open symbols) and 4 (filled symbols) (B). n =3.

**Supplementary Figure 4**

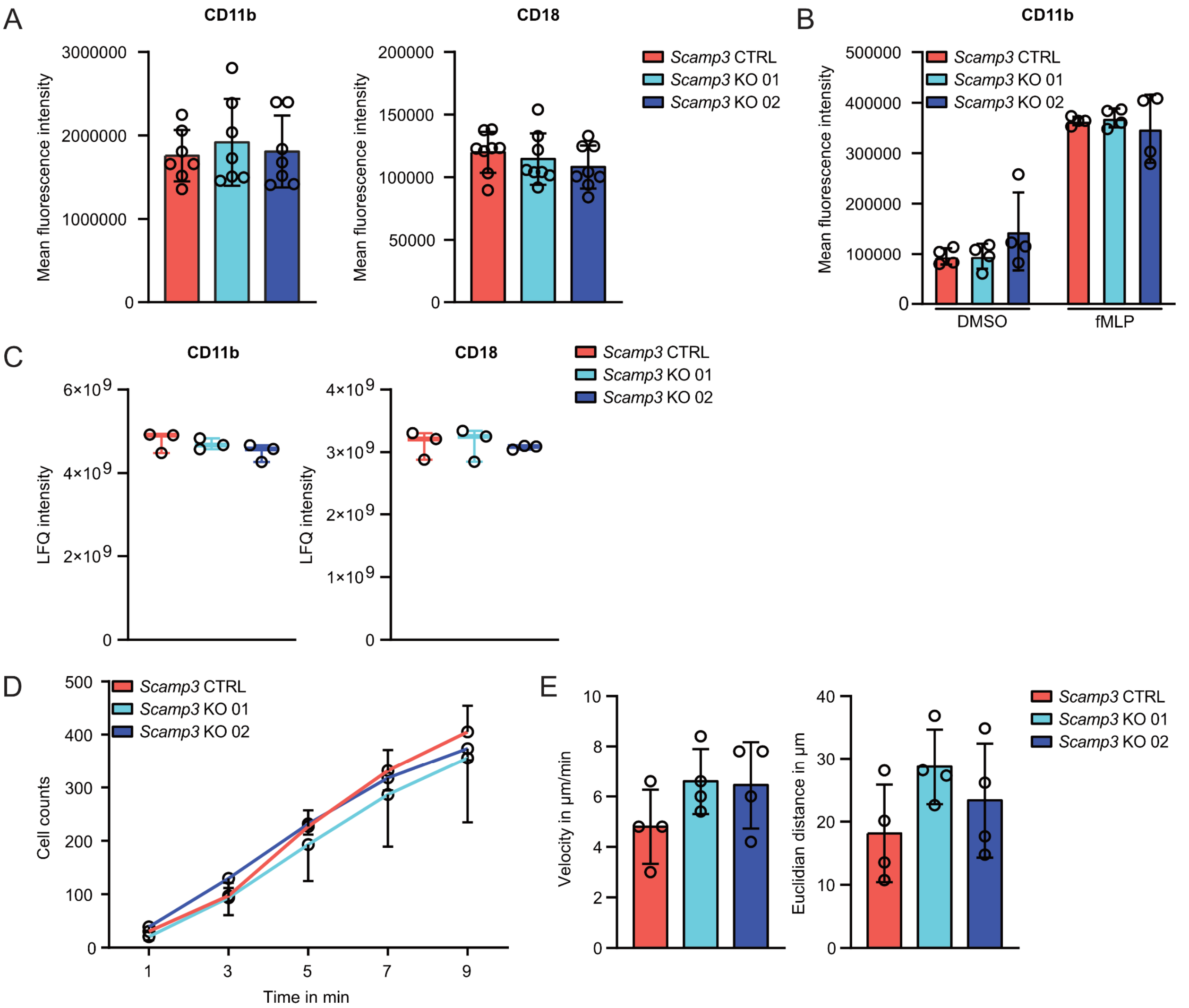


**Supplementary Figure 4.** Expression of $\beta_2$ integrin subunits and neutrophil recruitment of *Scamp3* CTRL and KO dHoxb8 cells. (A) Cell surface expression of CD11b and CD18 was analyzed by flow cytometry using fluorescently labeled antibodies. Mean fluorescence intensity is shown. n≥7. (B) Surface levels of CD11b after stimulation with fMLP (10 µM) or DMSO (0.15%) as control was analyzed by flow cytometry. n=4. (C) Total expression levels of CD11b and CD18 were analyzed by mass spectrometry. n=3. (D) Cell counts of adherent *Scamp3* control (CTRL) and KO dHoxb8 cells in microfluidic chambers coated with rmP-selectin (5 µg/ml), rmICAM-1 (3 µg/ml), and rmCXCL-1 (5 µg/ml) over 9 min of constant flow (1 dyne/cm²). n=6. (E) Velocity (left) and Euclidian distance (right) of *Scamp3* control (CTRL) and KO1 and KO dHoxb8 cells migrating in microfluidic chambers coated with rmICAM-1 (3 µg/ml) and rmCXCL-1 (5 µg/ml) under constant flow (1 dyne/cm²). n=4.

### Supplementary Figure 5

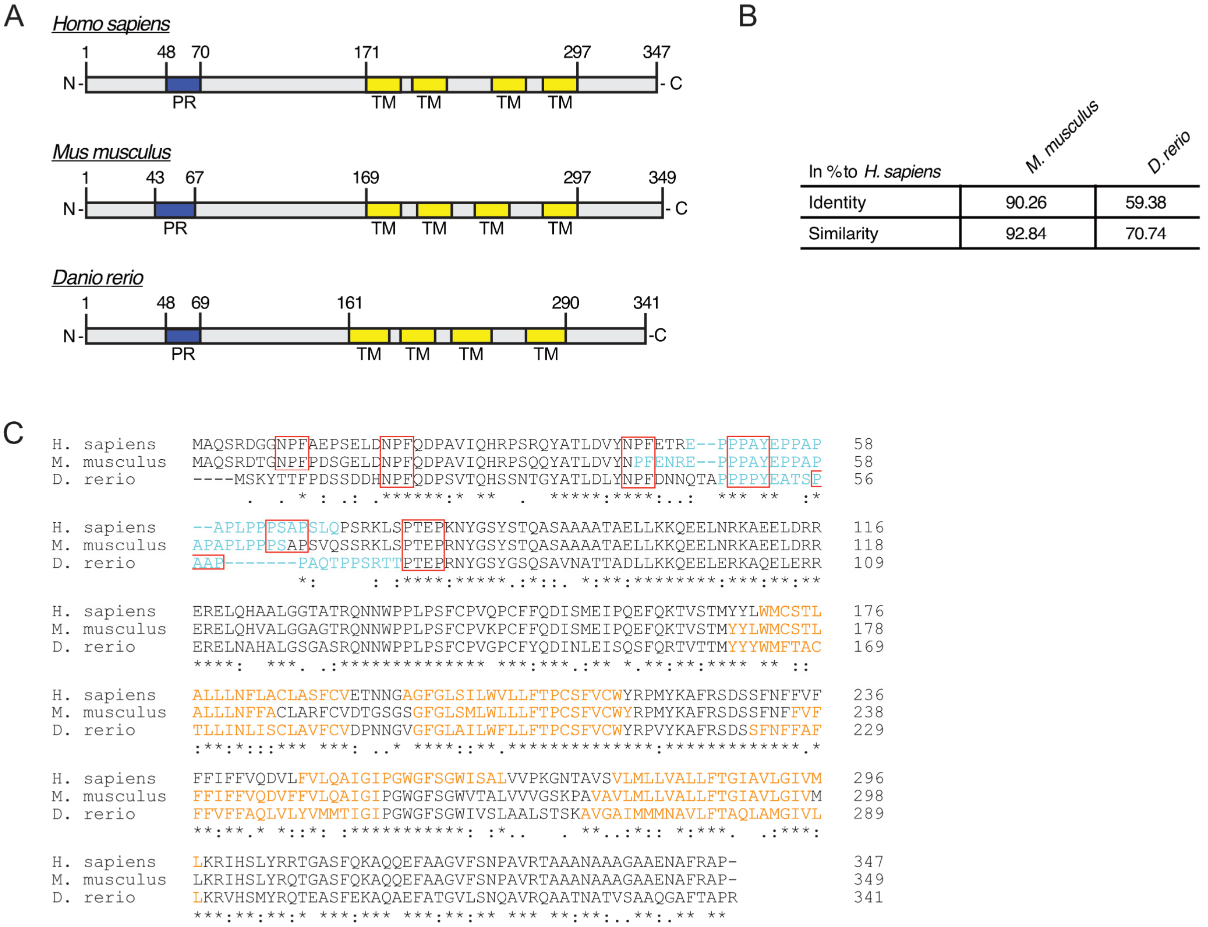


**Supplementary Figure 5.** Human, murine and zebrafish SCAMP3 protein and host defense functions in *scamp3* WT and KO zebrafish larvae. (A) Schematic of SCAMP3 protein from *Homo sapiens, Mus musculus* and *Danio rerio.* PR, proline rich region. TM, transmembrane domain. Numbers indicate amino acid positions. (B-C) Identity and similarity of human SCAMP3 to murine and zebrafish SCAMP3. (C) Sequence alignment of human, murine and zebrafish SCAMP3 using ClustalOmega. Identical (*), highly similar (:) and weakly similar (.) amino acids. Proline rich regions, blue. Transmembrane domain, orange. Conserved NPF, PPxY, PSAP and PTEP motifs, red boxes.
